# Objective versus Self-Reported Energy Intake Changes During Low-Carbohydrate and Low-Fat Diets

**DOI:** 10.1101/421321

**Authors:** Juen Guo, Jennifer L. Robinson, Christopher Gardner, Kevin D. Hall

**Affiliations:** National Institute of Diabetes and Digestive and Kidney Diseases; Stanford University

## Abstract

**Objective:** To examine objective versus self-reported energy intake changes (ΔEI) during a 12-month diet intervention.

**Methods:** We calculated ΔEI in subjects who participated in a 1-year randomized low-carbohydrate versus low-fat diet trial using repeated body weight measurements as inputs to an objective mathematical model (ΔEI_Model_) and compared these values with self-reported energy intake changes assessed by repeated 24-hr recalls (ΔEI_24hrRecall_).

**Results:** ΔEI_24hrRecall_ indicated a relatively persistent state of calorie restriction ≥500 kcal/d throughout the year with no significant differences between diets. ΔEI_Model_ demonstrated large early decreases in calorie intake >800 kcal/d followed by an exponential return to approximately 100 kcal/d below baseline at the end of the year. The low-carbohydrate diet resulted in ΔEI_Model_ that was 162±53 kcal/d lower than the low-fat diet over the first 3 months (p=0.002), but no significant diet differences were found at later times. Weight loss at 12 months was significantly related to ΔEI_Model_ at all time intervals for both diets (p<0.0001).

**Conclusions:** Self-reported measurements of ΔEI were inaccurate. Model-based calculations of ΔEI found that instructions to follow the low-carbohydrate diet resulted in greater calorie restriction than the low-fat diet in the early phases of the intervention, but these diet differences were not sustained.

**What is already known about this subject?:**

- Diet assessments that rely on self-report, such as 24hr dietary recall, are known to underestimate actual energy intake as measured by doubly labeled water. However, it is possible that repeated self-reported measurements could accurately detect changes in energy intake over time if the absolute bias of self-reported of measurements is approximately constant for each subject.

**What this study adds:**

- We compared energy intake changes measured using repeated 24hr dietary recall measurements collected over the course of the 1-year Diet Intervention Examining The Factors Interacting with Treatment Success (DIETFITS) trial versus energy intake changes calculated using repeated body weight measurements as inputs to a validated mathematical model.
- Whereas self-reported measurements indicated a relatively persistent state of calorie restriction, objective model-based measurements demonstrated a large early calorie restriction followed by an exponential rise in energy intake towards the pre-intervention baseline.
- Model-based calculations, but not self-reported measurements, found that low-carbohydrate diets led to significantly greater early decreases in energy intake compared to low-fat diets, but long-term energy intake changes were not significantly different.

## Introduction

Diet assessment instruments that rely on self-report, such as 24-hr recall, are known to substantially underestimate energy intake (1). However, repeated self-reported measurements could possibly track changes in energy intake accurately if the measurement bias is roughly constant for each subject. For example, if a person habitually eats a weight maintenance diet of 2500 kcal/d then their 24-hr recall might under-report eating only 1900 kcal/d. If they consistently underestimated their energy intake, then after starting a weight loss diet program they might report eating 1400 kcal/d whereas they actually consumed 2000 kcal/d. Their reported absolute energy intake would still be 600 kcal/d too low, but the self-reported change in energy intake of 500 kcal/d would be accurate. It is presently unknown whether people can accurately report changes in energy intake during a weight loss intervention.

We recently validated an objective mathematical method for calculating energy intake changes over time using only information about age, sex, height, and repeated measurements of body weight (2). Here, we applied this method to data from the Diet Intervention Examining The Factors Interacting with Treatment Success (DIETFIITS) randomized weight loss trial (3) and compared the model-calculated energy intake changes with self-reported values determined by repeated 24hr recalls.

## Methods

We used data from 414 subjects in the DIETFITS study (209 subjects randomized to the low-carbohydrate diet and 205 subjects randomized to the low-fat diet) with complete body weight data at all clinic visits. As previously described (3), weight was measured by digital scale at the Stanford Clinical Translational Research Unit and self-reported dietary intake was assessed using 3 unannounced 24-hour multiple-pass recall interviews (2 on weekdays and 1 on a weekend day) administered before the intervention and again after approximately 3, 6 and 12 months. Self-reported body weight was also recorded when subjects participated in the 22 instructional sessions over the course of the year. Participants were randomized to the low-carbohydrate or low-fat diet groups and were instructed reduce intake of total fat or digestible carbohydrates to 20 g/d during the first 8 weeks and then slowly add fats or carbohydrates back to their diets in increments of 5 to 15 g/d per week until they reached the lowest level of intake they believed could be maintained indefinitely.

As previously described (2), we used a linearized mathematical model of body weight dynamics solved for the average change in energy intake as compared to a weight-maintaining baseline diet, ΔEI_Model_, as a function of body weight and its rate of change as follows:

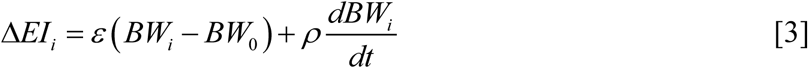

We used the initial age, sex, and height, along with an assumed initial free-living physical activity level (PAL) ∼1.6, to calculate the linearized model parameters for each subject (mean ± SE): ρ = 10036 ± 21 kcal/kg was the effective energy density associated with the change in body weight and ε = 23 ± 0.05 kcal/kg/d was the change in energy expenditure per unit body weight change. The change of mean body weight versus baseline over each interval, (*BW*_*i*_ – *BW*_0_), and the moving average of the measured body weight time course was used to calculate the rate of change of body weight over each interval, *dBW*_*i*_/*dt*. The interval length was *t* = *(N-1)*T*, where *N* was the number of body weight measurements per interval and *T* was the number of days between measurements. When clinic weights were used, N=2 for all periods and T=90 days for the first and second 3-month periods and T=180 days for the final 6 months. When self-reported weights were used, we specified the interval lengths of t=30 days, t=60 days, and t=90 days to calculate the average ΔEI_Model_ and the values for N and T were calculated using the available data on each subject in the corresponding time interval. In the figures, ΔEI_Model_ values were plotted at the midpoint time of the averaging interval.

Statistical analysis was performed using a paired, two-sided t-test with significance declared at the p<0.05 threshold. The data are reported as mean±SE.

## Results

**Figure 1A** shows the mean weight changes measured at the clinic visits and **Figure 1B** illustrates the model-based measurements as well as the self-reported measurements of energy intake change. After 3 months of the intervention, ΔEI_24hrRecall_ =-641±31 kcal/d which was significantly lower than ΔEI_24hrRecall_=-547±32 kcal/d at 6 months (p<0.0001). At 12 months, ΔEI_24hrRecall_=-500±31 kcal/d and was similar to the value at 6 months (p=0.05) indicating a relatively persistent and substantial reduction of energy intake.

**Figure 1.**
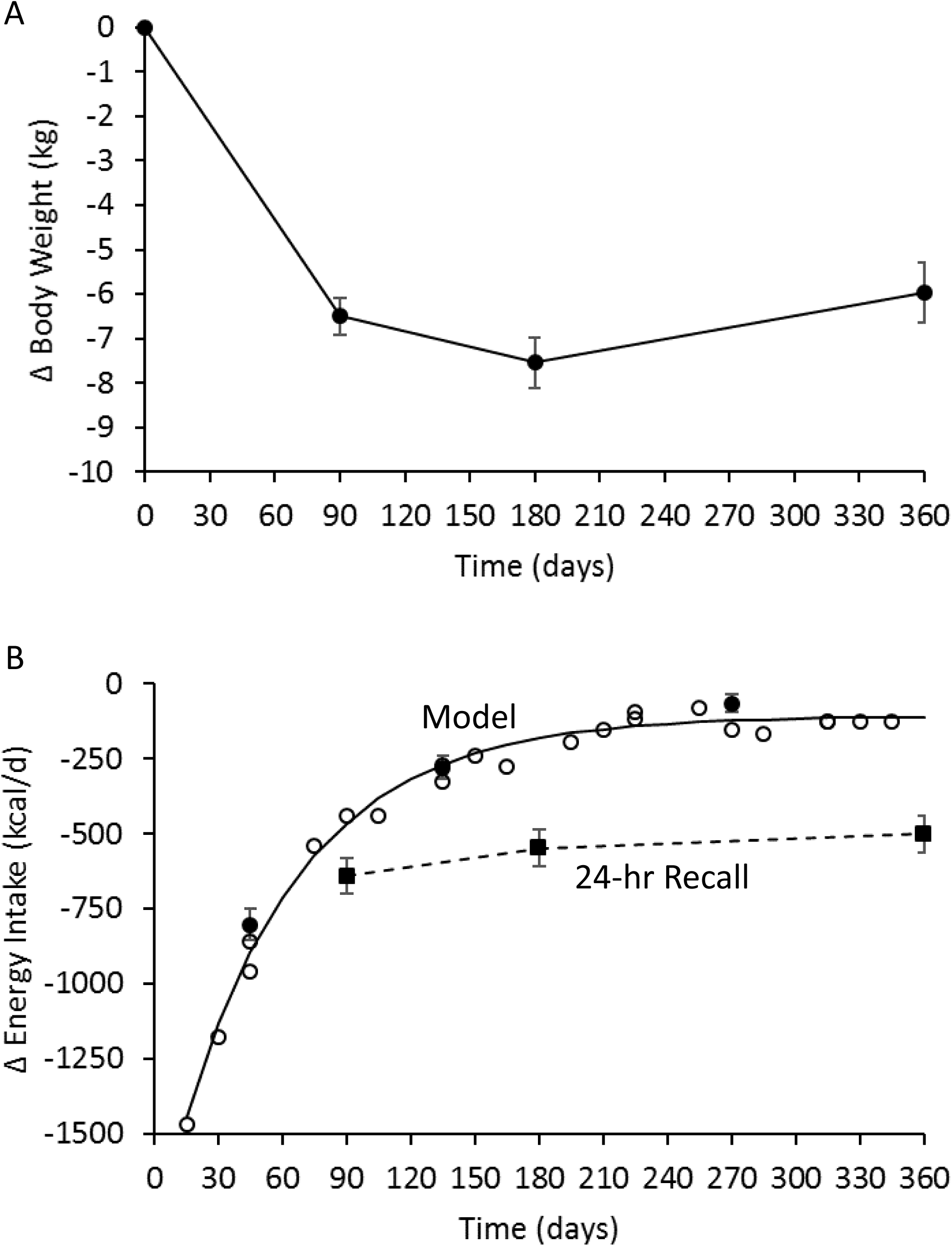
A) Mean body weight changes (⋅) measured during the DIETFITS trial clinic visits for all 414 subjects with complete clinic weight data. B) Mean self-reported energy intake changes (▪) indicated a relatively persistent reduction in energy intake whereas the model-based measurements (○ from self-reported weights and ⋅ from clinic weights) followed an exponential time course (solid curve). Error bars indicate 95% CI.

In contrast, the model-based calculations demonstrated that energy intake changes followed an exponential time course shown in Figure 1B. Using the clinic weights, ΔEI_Model_ was −804 ±27 kcal/d over the first 3 months. Over the next 3 months, ΔEI_Model_=-279 ±20 kcal/d indicating a substantial relaxation of calorie restriction (p<0.0001) which was again relaxed to ΔEI_Model_=-65 ±14 kcal/d between 6 and 12 months (p<0.0001).

**Figure 2A** shows the mean clinic weight changes in the low-carbohydrate and low-fat diet groups which were significantly different at 3 and 6 months, but not at 12 months. Self-reported energy intake was not significantly different between low-carbohydrate and low-fat diet groups at any time point (**Table 1**). However, model-based calculations using the clinic weights found that energy intake decreased over the first 3 months by 162±53 kcal/d more with the low-carbohydrate diet group as compared to the low-fat diet (p=0.002), but there were no significant differences at later times. Figure 2B shows that ΔEI_Model_ followed a similar exponential pattern regardless of diet, but the low-carbohydrate diet led to larger early reductions in calorie intake that were not sustained.

**Figure 2.**
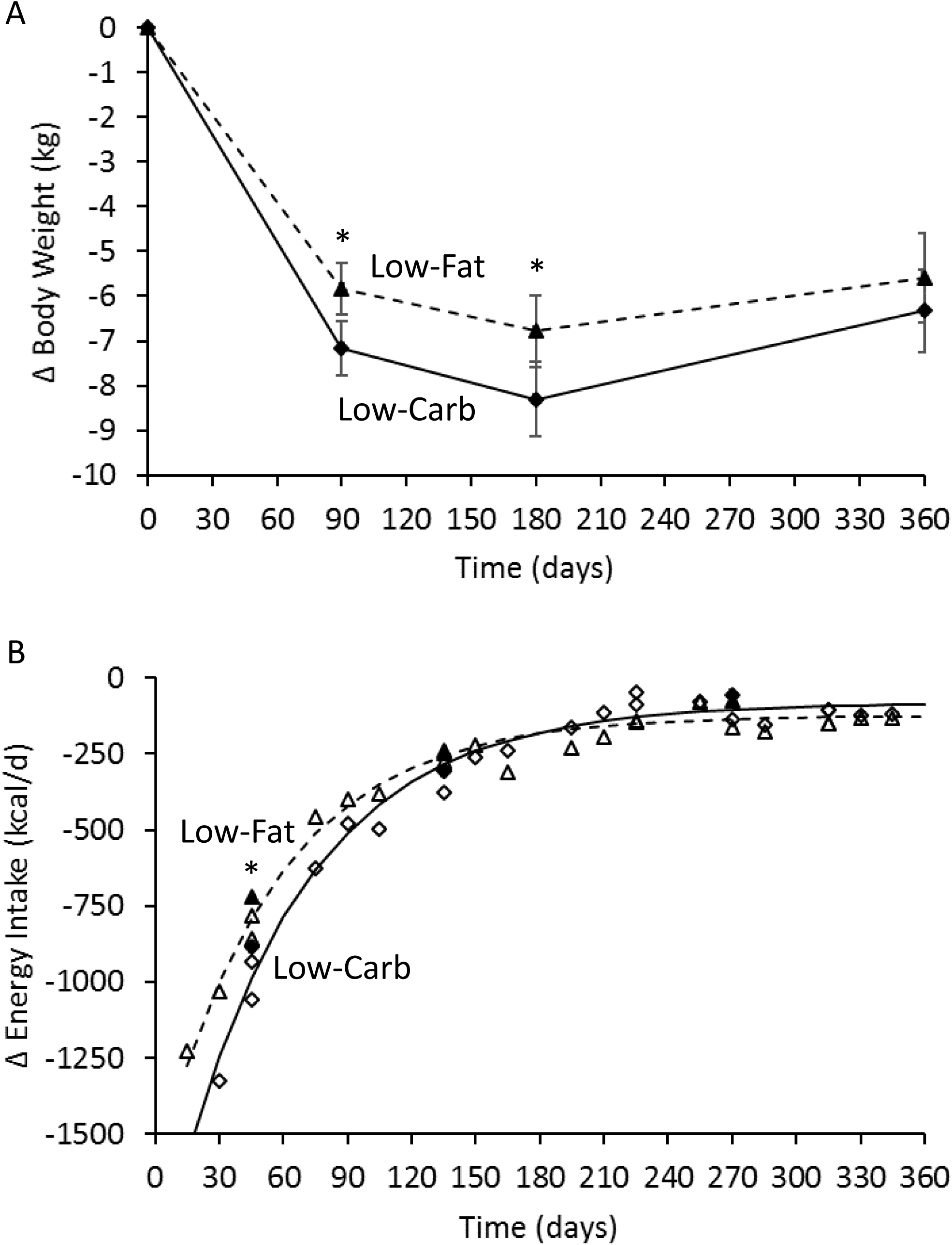
A) Mean body weight changes for the 209 subjects in the low-carbohydrate (♦) and the 205 subjects in the low-fat (▲) diet groups measured during the DIETFITS trial clinic visits. B) Mean model-based measurements of energy intake changes in the low-carbohydrate diet group (Δ from self-reported weights and ▲ from clinic weights) and the low-fat diet group (◊ from self-reported weights and ♦ from clinic weights) both followed an exponential time courses (solid curve and dashed curve for low-carbohydrate and low-fat diets, respectively). * indicates p<0.05 between diet groups and the error bars indicate 95% CI.

**Table 1.**

| **Variable** | **Both Diets**<br>**(N=414)** | **Low-**<br>**Carbohydrate**<br>**(LC; N=209)** | **Low-Fat**<br>**(LF; N=205)** | **P-value**<br>**LC vs LF** |
| --- | --- | --- | --- | --- |
| $\Delta BW_{3 \text{ months}}$ | -6.5±0.2 kg | -7.2±0.3 kg | -5.8±0.3 kg | 0.002 |
| $\Delta BW_{6 \text{ months}}$ | -7.6±0.3 kg | -8.3±0.4 kg | -6.8±0.4 kg | 0.01 |
| $\Delta BW_{12 \text{ months}}$ | -5.9±0.3 kg | -6.3±0.5 kg | -5.6±0.5 kg | 0.29 |
| 3 month $\Delta EI_{\text{Recall}}$ | -641±31 kcal/d | -628±43 kcal/d | -653±44 kcal/d | 0.68 |
| 6 month $\Delta EI_{\text{Recall}}$ | -547±32 kcal/d | -552±45 kcal/d | -542±45 kcal/d | 0.87 |
| 12 month $\Delta EI_{\text{Recall}}$ | -500±31 kcal/d | -532±44 kcal/d | -467±44 kcal/d | 0.30 |
| 0-3 month $\Delta EI_{\text{Model}}$ | -804±27 kcal/d | -884±39 kcal/d | -722±36 kcal/d | 0.002 |
| 3-6 month $\Delta EI_{\text{Model}}$ | -279±20 kcal/d | -307±29 kcal/d | -251±27 kcal/d | 0.16 |
| 6-12 month $\Delta EI_{\text{Model}}$ | -65±28 kcal/d | -56±18 kcal/d | -75±22 kcal/d | 0.49 |

**Figure 3** depicts individual 12 month weight change data for both the low-fat (left column) and low-carbohydrate (right column) diets as a function of the ΔEI_Model_ calculated using clinic weights averaged over the periods 6-12 months (panel A), 3-6 months (panel B), and 0-3 months (panel C). For the low-fat diet, weight loss at 12 months was correlated with ΔEI_Model_ averaged over 6-12 months (r=0.88; p<0.0001), 3-6 months (r=0.79; p<0.0001), and 0-3 months (r=0.70; p<0.0001). Weight change at 6 months was correlated with ΔEI_Model_ averaged over 3-6 months (r=0.88; p<0.0001), and 0-3 months (r=0.90; p<0.0001) and weight change at 3 months was correlated with ΔEI_Model_ averaged over 0-3 months (r=1; p<0.0001) (not shown). For the low-carbohydrate diet, weight loss at 12 months was correlated with ΔEI_Model_ averaged over 6-12 months (r=0.85; p<0.0001), 3-6 months (r=0.77; p<0.0001), and 0-3 months (r=0.70; p<0.0001). Weight change at 6 months was correlated with ΔEI_Model_ averaged over 3-6 months (r=0.85; p<0.0001), and 0-3 months (r=0.87; p<0.0001) and weight change at 3 months was correlated with ΔEI_Model_ averaged over 0-3 months (r=1; p<0.0001) (not shown). In contrast, ΔEI_24hrRecall_ was only weakly correlated with contemporaneous weight losses at 3-months (r=0.18; p=0.01) and 12-months (r=0.18; p=0.01) and only for the low-fat diet.

**Figure 3.**
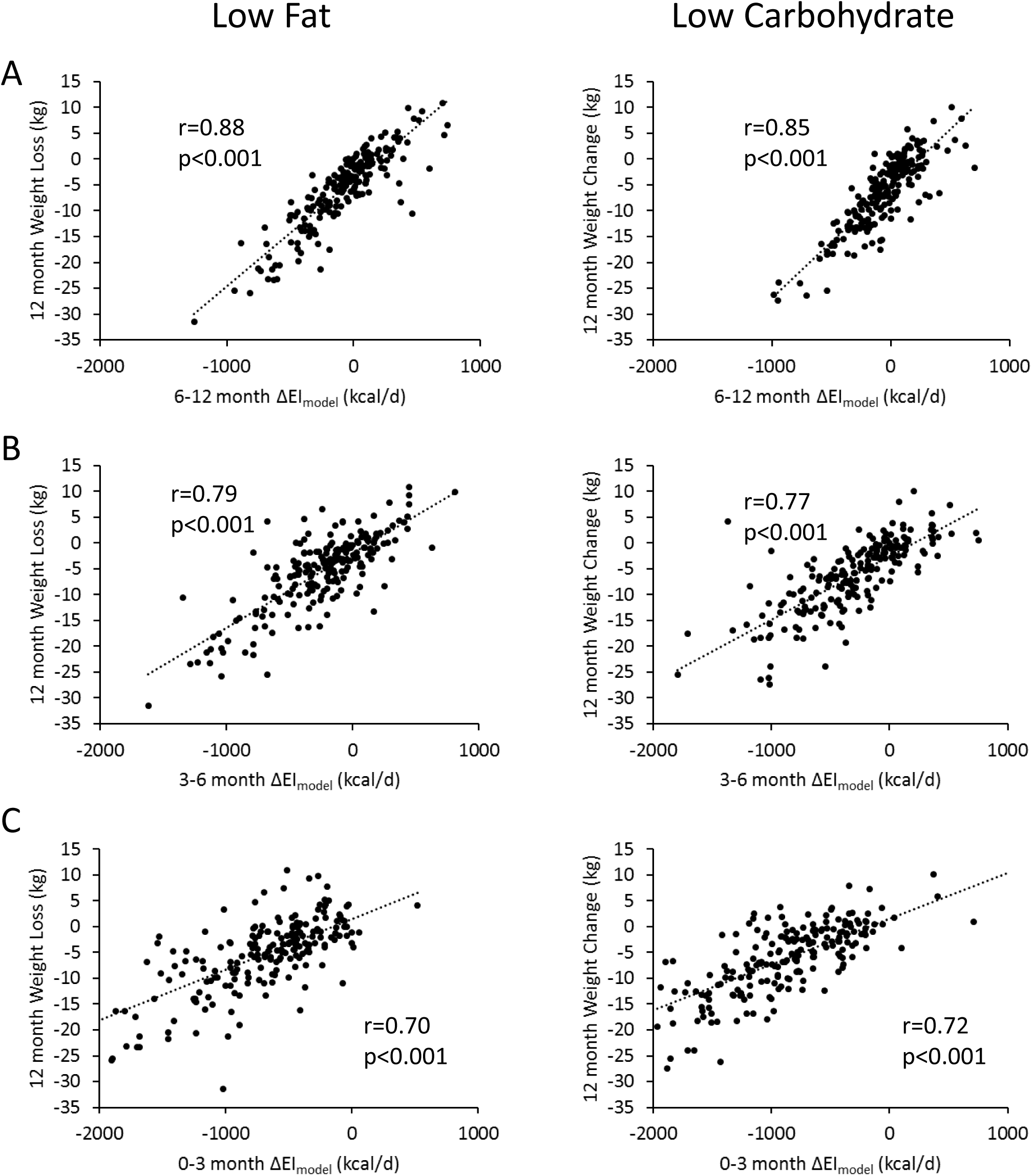
Individual weight changes at 12 months for subjects assigned to the low-fat diet (left column) and low-carbohydrate diet (right column) were significantly correlated with model-calculated changes in energy intake averaged over A) 6-12 months; B) 3-6 months; and C) 0-3 months.

## Discussion

This study demonstrates that the energy intake bias calculated by self-reported 24hr recall was not constant over time in subjects participating in a low-fat versus low-carbohydrate diet intervention for weight loss. Rather, biases in self-reported energy intake become progressively larger such that early assessments of ΔEI_24hrRecall_ were closer to ΔEI_Model_ as compared to later measurements. Whereas the ΔEI_24hrRecall_ measurements suggested a relatively persistent change in energy intake over time, the calculated average ΔEI_Model_ exhibited a large initial reduction in energy intake that exponentially decayed towards baseline over time. The low-carbohydrate diet resulted in significantly greater early reductions in energy intake, with correspondingly greater early weight losses as compared to the low-fat diet, but these diet differences were not sustained.

The exponential pattern of model-calculated energy intake with both diets is consistent with subjects exerting a relatively persistent effort to adhere to the diet intervention in the face of progressively increasing appetite in proportion to lost weight (4, 5). The self-reported reductions in energy intake were likely more representative of their persistent dieting efforts rather than indicating substantial sustained reductions in average energy intake.

During the initial stages of the low-carbohydrate diet, participants were instructed to reduce digestible carbohydrates to <20 g/d for the first 8 weeks and slowly add back carbohydrates to the minimum sustainable level (3). In this early time period, there was a greater reduction in model-calculated energy intake which is consistent with the suggestion that such very low carbohydrate diets suppress appetite by inducing nutritional ketosis (6). But such short-term reductions in appetite did not result in sustained reductions in energy intake with the low-carbohydrate diet and long-term weight loss was not significantly different between the diets.

At the end of the 12-month DIETFITS trial, there was a large interindividual variability in weight loss that was associated with the model-calculated energy intake changes at all stages of the intervention. Due to the long time-scale for human body weight to equilibrate to a constant energy intake (7), weight changes over periods of less than a few years are expected to be related to not only current energy intake, but the history of intake changes in the past year or more. Here, we observed that much of the 12-month weight loss variability was associated with energy intake changes occurring in the first few months as well as at later time points. Thus, studies designed to understand weight loss variability need to account for the dynamic nature of human weight loss.

The major limitation of this study was that we did not use doubly labeled water to measure free-living energy intake changes by the gold-standard intake-balance method (8). However, our mathematical method has been validated against the intake-balance method in a two year human calorie restriction study (2) that also exhibited a consistent exponential pattern of energy intake changes over time (9). However, this previous calorie restriction study did not compare different diets and did not include subjects with obesity (10), so we cannot be certain that the model-based calculations of energy intake were valid in the present study population.

In summary, repeated self-reported measurements of energy intake changes during the DIETFITS weight loss intervention were not accurate. Model-based calculations demonstrated an exponential pattern of energy intake change whereby large early calorie reductions decay back towards baseline over time. Instructions to adhere to a low-carbohydrate diet resulted in greater calorie restriction compared to a low-fat diet in the early phases of the DIETFITS intervention, but these diet differences were not sustained.

## Acknowledgements

KDH and JG were supported by the Intramural Research Program of the National Institutes of Health, National Institute of Diabetes and Digestive and Kidney Diseases. JLR and CG were supported by Grant 1R01DK091831 from the NIDDK, Grant 1K12GM088033 from the NIH (CTSA) and the Nutrition Science Initiative.

